# Alignment-driven Cross-Graph Modeling for 3D RNA Inverse Folding

**DOI:** 10.1101/2025.05.23.655885

**Authors:** Shengfan Wang, Hong-Bin Shen, Xiaoyong Pan

**Author notes:** Correspondence to: Xiaoyong Pan < >. Proceedings of the 41 ^st^ International Conference on Machine Learning, Vancouver, Canada. PMLR 267, 2025. Copyright 2025 by the author(s).

## Abstract

The 3D RNA inverse folding task aims to design RNA sequences that can fold into a target tertiary structure. This task is complicated by the inherent structural complexity and flexibility of RNA molecules. Considering that conserved RNA structures often exhibit sequence similarity, we present AlignIF, a pipeline designed to generate RNA sequences conditioned on the target RNA 3D structure and its multiple structure alignment (MStA), explicitly incorporating structural evolutionary conservation. AlignIF introduces a novel MStA backbone encoder designed to model multiple geometric graphs derived from MStA, which can better capture meaningful information from MStA structures. Additionally, AlignIF uses a random-ordered autoregressive sequence decoder to generate candidate RNA sequences. In our constructed benchmark dataset, AlignIF achieves a perplexity of 2.32 and a recovery rate of 57. 08%, respectively, a relative improvement of 17.8% and 11.8% over the state-of-the-art methods. In addition, AlignIF excels in diversity and ranking capability. We further demonstrate the added value of the incorporated MStA structures along with the proposed encoder architecture.

## 1. Introduction

RNA molecules are essential for cellular function. They serve as intermediates in protein synthesis and key regulators of gene expression, catalysis, and molecular scaffolding (Guil & Esteller, 2015; Kushwaha et al., 2016; Hentze et al., 2018). Recently, non-coding RNAs (ncRNAs) have attracted significant attention for their pivotal roles in regulating various cellular processes, such as gene expression, RNA modifications and splicing, genome stability, and regulation of cellular processes (Lakhotia et al., 2020; Beermann et al., 2016). While much research has focused on coding RNAs (Zhang et al., 2023; Metkar et al., 2024; Gong et al., 2023), the complex folding and functional mechanisms of ncRNAs—such as aptamers, riboswitches, and ribozymes—remain poorly understood. These ncRNAs’ three-dimensional (3D) structure is closely linked to their biological functions, playing a crucial role in their activity (Fu, 2014). Understanding these 3D conformations has become a key challenge in the field of RNA biology.

Recently, 3D RNA inverse folding has emerged as a powerful tool for addressing this challenge (Leman et al., 2020; Joshi et al., 2023; Tan et al., 2024; Huang et al., 2024; Wong et al., 2024). This approach focuses on designing RNA sequences that can fold into specific 3D structures, providing a transformative framework for studying and engineering ncRNAs. By enabling the design of RNA sequences that fold into desired 3D conformations, it holds significant potential for advancing our understanding of ncRNA functions.

However, RNA conformations exhibit significant dynamism and flexibility, with a single RNA sequence capable of folding into multiple distinct structures (Spitale & Incarnato, 2023). Furthermore, RNA structures are inherently complex and unstable, and the availability of experimentally determined 3D structural data remains limited (Zhang et al., 2022b). As a result, capturing the complete set of possible structures for a given RNA sequence is challenging. This limitation creates a substantial gap between the known sequence-structure joint distribution and the true distribution, which may lead to severe overfitting. Consequently, effectively utilizing the available structures to enhance sequence design remains a critical and unresolved challenge.

While gRNAde (Joshi et al., 2023) leverages multiple conformational states of the same RNA (i.e., different structures folded from the same sequence) to generate candidate sequences, its approach has notable limitations. It assumes prior knowledge of the true sequence, which is typically impractical prior to inverse folding. Furthermore, in most cases, only a single RNA conformational state is accessible, restricting its applicability.

To address this challenge, we propose AlignIF, a novel method for designing RNA sequences based on a target structure and its homologous structures. AlignIF draws inspiration from structure prediction, where multiple sequence alignments (MSA) are constructed from a single input sequence to capture evolutionary information, thereby improving the accuracy of 3D structure prediction (Shen et al., 2024; Abramson et al., 2024). For the inverse problem of RNA sequence design based on a target structure, we introduce multiple structure alignments (MStA), which capture evolutionary conservation by aligning similar structures to the target structure at the nucleotide level. MStA can be constructed using US-align (Zhang et al., 2022a) and is searched from experimentally determined database (PDB).

AlignIF consists of an MStA backbone encoder and an autoregressive sequence decoder. The encoder processes geometric features of individual structures through message passing and integrates information across structures using the self-attention mechanism. The decoder leverages the learned geometric representations of the target structure to generate RNA sequences that can fold into the desired conformation.

Our contributions are listed as follows:

- We propose AlignIF, a simple and effective method for RNA sequence design based on the target backbone structure and integrated MStA structures.
- We evaluate AlignIF on a curated RNA inverse folding benchmark, assessing the native perplexity, native recovery rate, diversity, and ranking capability. AlignIF achieves significant improvements over existing 3D RNA inverse folding methods, including RDesign (Tan et al., 2024), RiboDiffusion (Huang et al., 2024), gRNAde (Joshi et al., 2023), and RhoDesign (Wong et al., 2024).
- We conduct an in-depth analysis of the contribution brought by the integrated MStA structures, highlighting their impact on the overall performance. Additionally, we evaluate the effectiveness of the proposed architecture in addressing the specific challenges of the RNA design task.

## 2. Related Work

### RNA structure prediction

RNA structure prediction is to determine a molecule’s two or three-dimensional (2D or 3D) structure from its sequence, essentially the reverse of inverse folding. Early approaches, such as Vienna RNAfold (Lorenz et al., 2011) and MFold (Zuker, 2003), concentrated on predicting RNA secondary structures by minimizing free energy. For RNA tertiary structure prediction, methods like SimRNA (Boniecki et al., 2016) utilized similar energy minimization techniques to model 3D conformations. Fragment assembly approaches, including FarFar (Das & Baker, 2007) and its successor FarFar2 (Watkins et al., 2020), were also developed to predict RNA 3D structures.

In recent years, deep learning has significantly advanced structure prediction. For RNA secondary structure prediction, tools such as SPOT-RNA (Singh et al., 2019; 2021), UFold (Fu et al., 2022), and RNA-MSM (Zhang et al., 2024) have emerged, demonstrating an improved accuracy and efficiency. For RNA tertiary structure prediction, models like trRosettaRNA (Wang et al., 2023), DRFold (Li et al., 2023), AlphaFold3 (Abramson et al., 2024), and RhoFold+ (Shen et al., 2024) are gaining prominence. Most of these methods harness multiple sequence alignments (MSA) to capture evolutionary conservation, a crucial factor in improving the predictive accuracy.

### RNA sequence design

RNA sequence design has been approached using various methodologies, which can be classified into unconditional design and conditional design.

Unconditional RNA sequence design focuses on generating sequences that reflect the native distribution of RNA. Methods such as RaptGen (Iwano et al., 2022) and RfamGen (Sumi et al., 2024) leverage Variational Autoencoders (VAEs) to map the sequence space into a latent representation, enabling the sampling of sequences that capture the underlying distribution of native RNAs.

For conditional RNA sequence design, the goal is to generate RNA sequences that satisfy specific properties or constraints. AptaDiff (Wang et al., 2024) integrates a Variational Autoencoder (VAE) with Discrete Denoising Diffusion Probabilistic Models (D3PM) to create sequences tailored to a given motif. LEARNA (Runge et al., 2018) and libLEARNA (Runge et al., 2023) leverage reinforcement learning to optimize policies that sample sequences closely resembling a specified target secondary structure. RNA-DCGen (Shahgir et al., 2024) employs an RNA language model to iteratively generate sequences that align progressively with a given RNA distance map and secondary structure. RNAFlow (Nori & Jin, 2024) adopts flow matching, combining inverse folding and structure prediction, to co-generate RNA sequences and structures that bind to a specific protein. Additionally, methods for 3D RNA inverse folding (Tan et al., 2024; Huang et al., 2024; Joshi et al., 2023; Wong et al., 2024) represent a category of conditional design focusing on sequence generation constrained by a fixed backbone structure.

## 3. Methods

This section introduces AlignIF, a novel RNA design method based on tertiary backbone structures and their corresponding MStA representations. AlignIF consists of two main components: (i) an MStA backbone encoder, which takes a given RNA structure and its corresponding MStA as input, and (ii) a sequence decoder that generates RNA sequences capable of folding into the target structure in a random order, rather than a fixed 5’ to 3’ direction. An overview of the AlignIF pipeline is provided in Figure 1. The key contributions of this work is the development of the MStA backbone encoder, which models multiple geometric graphs derived from MStA structures. This encoder integrates aligned features of nodes and edges at each layer, enabling it to capture structural evolutionary conservation effectively.

**Figure 1.**
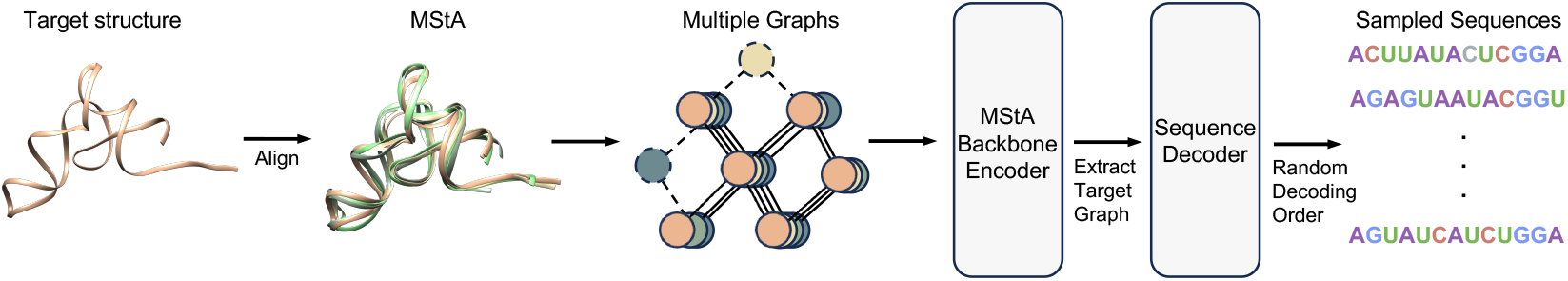
The overview of our proposed AlignIF.

### Representing MStA as Multiple Graphs

#### Node And edge featurization

Each RNA structure is represented as a graph *𝒢* = (*𝒱, ε*). For a structure with *N* nucleotides, the node set *𝒢* = *{****v***_1_, …, ***v***_*N*_ *}* encodes the features of individual residues, capturing inter-residue properties. The edge set *ε* = {***e***_*ij*_} _*j∈풩* (*i,r*)_ represents relationships between node *i* and its neighbors, defined as nodes within a spatial distance of *r*. Here, 𝒩 (*i, r*) denotes the neighbors of node *i* within this threshold. Each nucleotide’s spatial location is defined by the 3D coordinates of its C3’ atom. To effectively model the backbone structure, we construct comprehensive geometric representations for both nodes and edges. The specific features and their corresponding illustrations are detailed in Table 1.

**Table 1.** The feature construction of the graph from RNA tertiary structure.

| LEVEL | FEATURE | ILLUSTRATION |
| --- | --- | --- |
| NODE | DIHEDRAL ANGLE | $\{\sin, \cos\} \times \{\alpha_i, \beta_i, \gamma_i, \delta_i, \varepsilon_i, \xi_i, \eta_i, \theta_i\}$ |
| | DIRECTION | $Q_i^T \frac{P_i - C3'_i}{\ P_i - C3'_i\ }, Q_i^T \frac{N_i - C3'_i}{\ N_i - C3'_i\ }$ |
| | ORIENTATION | $q(Q_i^T Q_{i-1}), q(Q_i^T Q_{i+1})$ |
| EDGE | DISTANCE | $\text{RBF} \times \{\ P_j - C3'_i\ , \ C3'_j - C3'_i\ , \ N_j - C3'_i\ , \ N_j - N_i\ \}$ |
| | ORIENTATION | $q(Q_i^T Q_j)$ |

#### Local coordinate system

Ensuring that RNA representation remains invariant to global rotations and translations is essential, as these transformations do not alter the underlying sequence. To address this, we introduce a local coordinate system, denoted as ***Q*** for each nucleotide. Unlike RDesign (Tan et al., 2024), which defines the local coordinate system using three atoms from consecutive bases, our approach avoids the limitations posed by RNA chain flexibility. Such flexibility renders the traditional systems highly dynamic and inconsistent, complicating the extraction of invariant features. Furthermore, defining a coordinate system at the beginning and end of a sequence is impractical due to its reliance on the presence of three consecutive residues.

To address these limitations, we propose a modified local coordinate system based on the positions of the C2’, C3’, and C4’ atoms within the rigid sugar ring of each nucleotide, as shown in Figure 2c. This system ensures consistent invariant feature extraction. A detailed explanation of this system is provided in Appendix B.

**Figure 2.**
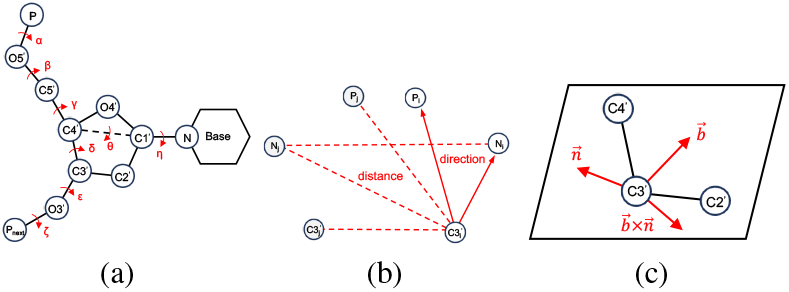
AlignIF extracts features from RNA backbone structures, including: (a) Seven natural dihedral angles and one pseudodihedral angle; (b) Distances and directions derived from the 3bead coarse-grained backbone model; (c) Local coordinate systems defined by the C2’, C3’, and C4’ atoms within the sugar ring.

#### Node-level features

We first calculate seven natural dihedral angles and a pseudo dihedral angle *θ*, as shown in Figure 2a. These dihedral angles are represented using sin and cos functions. Next, we compute the directions of the P and C3’ atoms relative to the C3’ atom in the local coordinate system ***Q***. Finally, two spatial rotation matrices, derived from the current system relative to the previous and next systems, are encoded using quaternion.

#### Edge-level features

We calculate the distances between the P, C3’, and N atoms (N1 for pyrimidines and N9 for purines) in neighboring nucleotides and the C3’ atom in the current nucleotide. Additionally, the N-N distance is included in the edge features. These distances are encoded using radial basis functions (RBFs). The orientation between two nucleotides is calculated in the same manner as the node orientation.

### 3.1.2 Multiple Graphs Construction from Structure Alignment

Starting with a given target structure, we obtain the MStA using the US-align tool (Zhang et al., 2022a), which accurately aligns RNA structures. US-align provides both a global alignment score (TM-score) and local alignment pairs. When the TM-score between two structures exceeds 0.45, it typically indicates that their folding patterns are globally similar and suggests potential similarities in their biological functions (Gong et al., 2019). Therefore, we select structures with a TM-score greater than 0.45 as members of MStA. These selected MStA structures are then converted into multiple graphs. Based on the local alignment pairs, only the aligned nodes and edges are retained in the graphs, while the unaligned parts are discarded, as shown by the dotted circles and lines in Figure 1. This approach ensures that dissimilar nodes and edges do not influence the model, preventing the generation of sequences that fold into incorrect structures.

### 3.2. Cross-Graph Modeling

When representing MStA structures as multiple graphs, 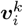 and 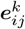 denote the *i*-th node and the edge between node *i and node j in the k-th structure, respectively. The AlignIF model processes multiple attributed graphs using the proposed MStA backbone encoder, invariant to graphs’ ordering. At the end of the encoder, the target graph is extracted for sequence decoding*.

#### MStA backbone encoder

We update geometric graphs through message passing, processing each graph independently. Subsequently, we integrate features across multiple graphs using a self-attention mechanism to jointly update the node and edge features of each graph, ensuring the permutation invariance is preserved. This approach is illustrated in Figure 3, which illustrates cross-graph modeling. Initially, we use message passing to update the features 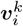for each

**Figure 3.**
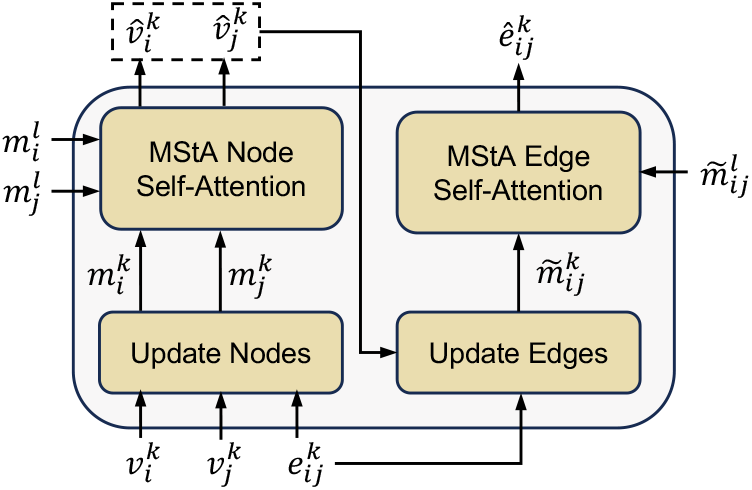
Update nodes and edges features across graphs.

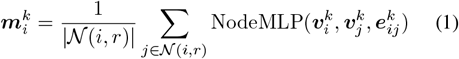

Next, the node feature 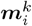 updated by message passing is processed using self-attention across all *K* structures:

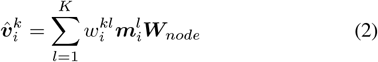

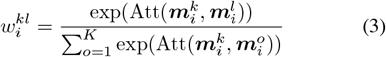

where ***W***_*node*_ is a learnable matrix used to project node features, 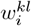 is the attention weight, and 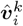 is the updated feature for node *i* in the *k*-th structure. In addition to updating node features, we similarly update edge features:

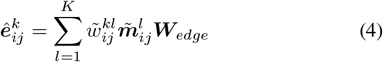

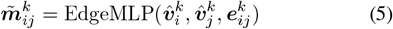

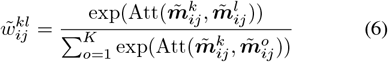

The proposed MStA backbone encoder iteratively updates features through message passing and an attention mechanism, facilitating cross-graph modeling. This design effectively integrates information from multiple graphs, allowing it to learn more robust and consistent representations.

#### Sequence decoder

The final encoder representations in AlignIF capture the MStA information while maintaining invariance to the structural permutations within MStA. To prevent the model from becoming confused by multiple structural inputs, we focus only on extracting the target graph, which consolidates information from other structures within MStA, as the input into the sequence decoder. In the decoder, the model autoregressively predicts each node’s probability of one of the four nucleotide identities (A, G, C, U). Following the approach of ProteinMPNN (Dauparas et al., 2022), we replace the conventional 5’ to 3’ terminal decoding order with an order-agnostic method that randomly samples from all possible permutations of base sequences. This flexible decoding strategy enables a more versatile RNA sequence design and reduces the risk of overfitting to specific sequence patterns. AlignIF is trained in a self-supervised manner to reconstruct the native sequence, minimizing the cross-entropy loss (with a label smoothing value of 0.1) between the predicted probabilities and the true nucleotide identity for each node. During training, teacher forcing (Williams & Zipser, 1989) is used, where the true previous token is fed into the decoder as input, rather than the model’s predictions, to facilitate more stable and faster convergence.

### 3.3. RNA Sequence Design Performance Metrics

We evaluate the quality of the designed RNA sequences using two key metrics: native perplexity and native recovery rate. Native perplexity quantifies the uncertainty of the native sequence, reflecting how well the model can generate the ground truth sequence. The native recovery rate measures the percentage of native nucleotides correctly recovered in the designed RNA sequences. In addition to perplexity and recovery, we assess the diversity of the generated RNA sequences. Diversity is calculated by comparing pairs of sequences from the generated set, measuring the proportion of nucleotide identity differences. Finally, we assess the ranking capability of AlignIF by correlating the perplexities of generated sequences with their corresponding native recovery rates.

## 4. Experiments

### 4.1 Implementation Details

The MStA backbone encoder and sequence decoder individually consist of 3 layers. Both the node and edge hidden features have a dimensionality of 128. The model is trained with a mini-batch size of 64 RNAs and a learning rate of 1e-4. Training is performed using a single NVIDIA GeForce RTX 4090 D GPU, with the Adam optimizer (Kingma, 2014). A label smoothing rate of 0.1, a dropout rate of 0.2, and an exponential moving average rate of 0.999 are applied during training. The distance threshold *r* to construct graph is set to 20 Å.

#### Benchmark dataset

The 3D RNA structure dataset is derived from the BGSU database (Leontis & Zirbel, 2012), which compiles all RNA-containing 3D structures from the Protein Data Bank (PDB) (Berman et al., 2000). We include only structures with a resolution of *≤* 4Å and filter out sequences containing more than 256 or fewer than 16 nucleotides. This results in a final dataset of 7,792 structures (before BGSU data release date: May 1, 2024). Additional dataset statistics, including rationale for the selected sequence length interval, sequence length distribution, and MStA size distribution, can be found in Appendix A.

To assess AlignIF’s generalization ability to novel RNAs, we cluster the 7,792 structures using a structural similarity threshold of TM-score *≥*0.45, yielding 467 clusters. This threshold ensures that structurally similar RNAs are grouped while guaranteeing that the MStA of each structure remains within the same cluster, thereby preventing information leakage. The dataset was then split into the training (7,458), validation (111), and test (223) sets, following an 8:1:1 ratio based on clusters.

#### Baseline Models

We evaluate AlignIF on the 3D RNA inverse design task by comparing it with recent baseline models, including RDesign (Tan et al., 2024), RiboDiffusion (Huang et al., 2024), gRNAde (Joshi et al., 2023), RhoDesign (Wong et al., 2024). To ensure a fair and reliable comparison, all baseline models are retrained using their default settings on the benchmark dataset.

### 4.2 RNA Design Quality Performance

This task is to design RNA sequences based on specified backbone structure. Our approach not only focuses on the given backbone structure but also integrates the corresponding MStA backbone structures, enhancing the design process.

We assess model performance on the test set, selecting those with the lowest loss on the validation set. To investigate the impact of RNA length on generative quality, we categorize the test set into three groups based on RNA length: Short (*≤*50 nucleotides), Medium (*>*50 but *≤*100 nucleotides), and Long (*>*100 nucleotides). We report native perplexity and recovery metrics for each group separately, as well as the overall performance on the test set. For autoregressive models (gRNAde, RhoDesign, and AlignIF) and the diffusion model (RiboDiffusion), which can generate multiple sequences for a given structure, we generate 20 sequences per structure (sampled at a temperature of 0.1 for the autoregressive models) and calculate the average values for both metrics. For RDesign, which predicts only a single sequence, we directly calculate the metrics for that sequence.

As shown in Table 2, the proposed AlignIF consistently improves native perplexity and recovery on the test set, with improvements of 5.11% and 0.5, respectively. RiboDiffusion is not included in the perplexity comparison, as it is a diffusion model and cannot compute perplexity. These gains are primarily attributed to AlignIF’s strong performance in the “Medium” and “Long” groups, indicating that AlignIF is capable of capturing more complex structural patterns.

**Table 2.** Comparison of native perplexity and recovery metrics on the benchmark dataset against other baseline models. The best and suboptimal results are highlighted in bold and underlined.

| MODEL | SHORT | PERPLEXITY ↓ |  | ALL | SHORT | RECOVERY % ↑ |  | ALL |
| --- | --- | --- | --- | --- | --- | --- | --- | --- |
|  |  | MEDIUM | LONG |  |  | MEDIUM | LONG |  |
| RDESIGN | 3.844 | 3.789 | 3.874 | 3.837 | 33.87 | 34.55 | 33.10 | 33.77 |
| RIBODIFFUSION | - | - | - | - | 38.18 | 42.43 | 37.97 | 39.81 |
| GRNADE | 2.867 | <u>2.705</u> | <u>2.964</u> | 2.849 | 38.18 | 43.77 | 41.02 | 41.50 |
| RHODESIGN | **2.304** | 2.819 | 3.073 | <u>2.824</u> | **59.72** | <u>55.38</u> | <u>46.05</u> | <u>51.97</u> |
| ALIGNIF | <u>2.627</u> | **2.085** | **2.418** | **2.324** | <u>51.79</u> | **61.82** | **55.41** | **57.08** |

### 4.3 Diversity Performance

We compare the trade-off between diversity and recovery across different models. For autoregressive models, sequences are generated at temperatures ranging from 1.0 to 0.1 in intervals of 0.1, with an additional generation at a temperature of 0.01. Each circle in Figure 4 represents a different temperature setting, with temperatures increasing from left to right. As the temperature increases, the diversity rises while the recovery decreases. For all models except RDesign, 20 sequences are generated for each temperature, and both diversity and average recovery are calculated. As shown in Figure 4, AlignIF achieves the highest diversity while maintaining the optimal recovery performance, indicating that AlignIF can simultaneously capture both conservative and flexible regions.

**Figure 4.**
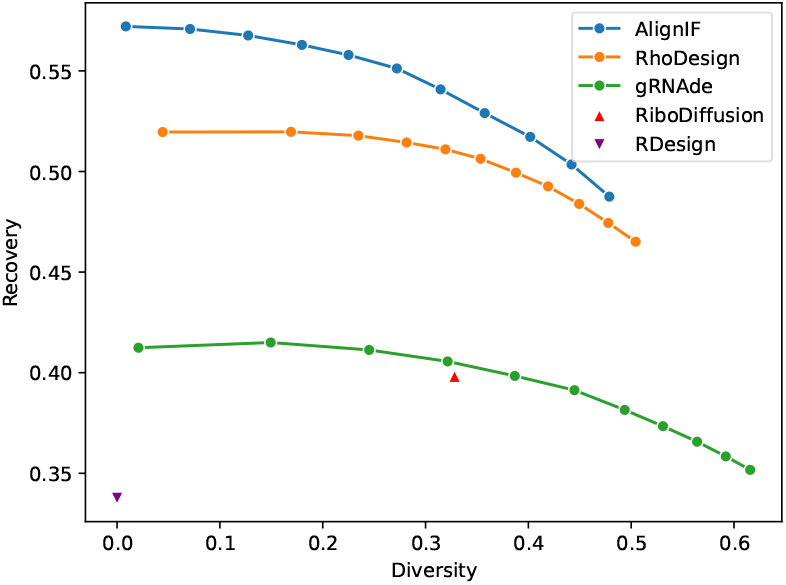
The relationship between the diversity and recovery, with each circle representing a different sampling temperature.

### 4.4 Ranking Capability Performance

Perplexity measures the level of “uncertainty” or “confusion” the model experiences when predicting the next token in a sequence. It does not require ground-truth sequences, which makes it useful for ranking sequences. The one exhibiting lower perplexity is considered better, reflecting the model’s higher confidence (lower uncertainty) in its prediction.

For each structure in the test set, we generate 20 sequences by sampling at temperatures of 0.1, 0.5, and 1.0. We then calculate both the Pearson correlation coefficient and the Spearman rank correlation coefficient to evaluate the relationship between perplexity and native recovery rate. We compare the results with those of other autoregressive baseline models, as shown in Table 3.

**Table 3.** Correlation between the perplexity and native recovery rate at three different temperatures.

| MODEL | PEARSON ↓ |  |  | SPEARMAN ↓ |  |  |
| --- | --- | --- | --- | --- | --- | --- |
|  | T = 0.1 | T = 0.5 | T = 1.0 | T = 0.1 | T = 0.5 | T = 1.0 |
| GRNADE | -0.47 | -0.55 | -0.54 | -0.53 | -0.61 | -0.55 |
| RHODESIGN | -0.60 | -0.64 | -0.65 | -0.57 | -0.64 | -0.65 |
| ALIGNIF | **-0.77** | **-0.78** | **-0.75** | **-0.83** | **-0.85** | **-0.79** |

In AlignIF, the perplexity shows a strong negative correlation with the native recovery rate. The Pearson and Spearman rank correlation coefficients for AlignIF surpass those of other autoregressive baseline models. These results underscore AlignIF’s efficiency in quickly ranking sequences, thereby facilitating the selection of candidates for experimental validation.

## 5. Analysis

### 5.1 Contribution of MStA Structures

RNA structures often undergo dynamic changes due to their inherent flexibility (Spitale & Incarnato, 2023). Multiple similar structures can capture these dynamic changes, particularly in conserved regions. To better capture this flexibility, we design sequences explicitly conditioned on both the target structure and its corresponding MStA structures.

To evaluate whether incorporating MStA structures enhances AlignIF’s design capability, we compared its performance across varying MStA depths (the maximum number of structures used from the MStA set) on the test set. The structures selected from the MStA set are greedily extracted based on their ranking TM-scores with the target structure, ensuring that only the most structurally similar models are included. As shown in Table 4, AlignIF consistently improves as the MStA depth increases. Notably, AlignIF is also capable of handling RNAs without MStA, due to the random masking of MStA during the training stage.

**Table 4.**
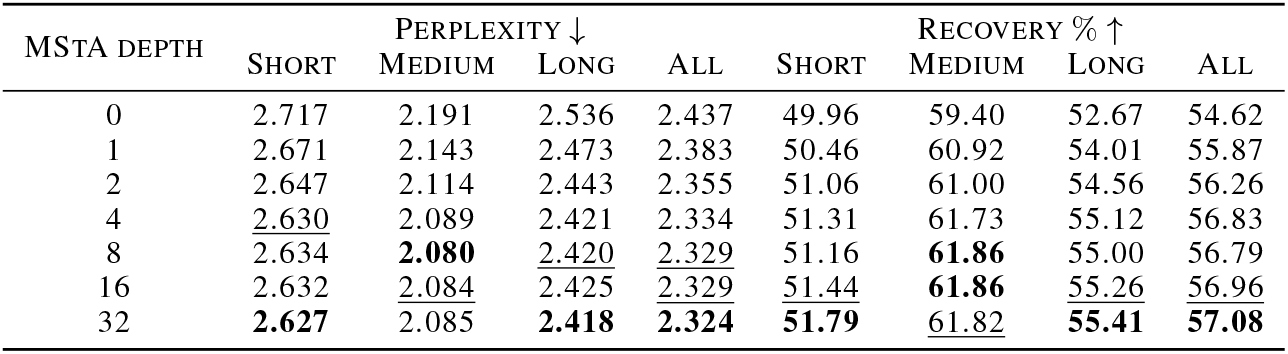
Perplexity and recovery metrics vary with the depth of MStA on the test set, with the best and suboptimal results highlighted in bold and underlined.

### 5.2 Contribution of the modules in AlignIF

To better understand the contributions of each module in AlignIF, we conduct an ablation study. In the MStA backbone encoder, we update nodes and edges concurrently across graphs. Previous study (Dauparas et al., 2022) have demonstrated the effectiveness of independent message-passing updates for nodes and edges within individual graphs. However, the impact of incorporating cross-graph node and edge updates remains to be further explored. Additionally, we modify the local coordinate system to achieve a more stable and invariant representation. Finally, we utilize a random-ordered autoregressive sequence decoder, which not only introduces more flexibility in the decoding process but also improves overall performance.

The ablation study includes the following variants: (i) AlignIF w/o node, which removes the cross-graph node update; (ii) AlignIF w/o edge, which excludes the cross-graph edge update; (iii) AlignIF w/o system, which uses the traditional local coordinate system; and (iv) AlignIF w/o random, which replaces the random-order decoder with a fixed-order decoder. The results in Table 5 indicate that all the incorporated modules are indispensable.

**Table 5.** Ablation study on the benchmark dataset.

| METHODS | PERPLEXITY ↓ | RECOVERY % ↑ |
| --- | --- | --- |
| ALIGNIF | **2.324** | **57.08** |
| W/O NODE | 2.462 | 56.35 |
| W/O EDGE | 2.428 | 54.52 |
| W/O SYSTEM | 2.437 | 54.21 |
| W/O RANDOM | 2.433 | 53.59 |

## 6. Conclusion and Limitations

In this study, we introduce AlignIF, a novel generative model for designing RNA sequences tailored to target backbone structures and their corresponding MStA graphs. AlignIF incorporates an MStA backbone encoder that iteratively updates node and edge representations across MStA graphs. Additionally, a random-ordered autoregressive decoder is employed to generate RNA sequences based on the learned target graph. Our approach consistently surpasses previous state-of-the-art 3D RNA inverse folding methods on a curated benchmark dataset, achieving superior perplexity, recovery, diversity, and ranking capabilities. However, AlignIF cannot generate RNAs with properties superior to those of existing RNA molecules. Additionally, our method is currently confined to in silico design, with wet-lab validation deferred for future studies.

### Impact Statement

This paper contributes to the design of sequences based on the target structure and its MStA structures, advancing the field of inverse folding. While our work has potential societal implications, we do not feel the need to specifically highlight them here.

## A. Benchmark dataset details

We present the histogram of sequence length distribution, as shown in Figure 5. The dataset displays a long-tailed distribution, with a high frequency of short sequences under 256 nucleotides. Most functional RNAs, such as aptamers, riboswitches, and ribozymes, predominantly fall within this range. RNA folding, which relies on nucleotide interactions, typically requires longer sequences to form stable secondary structures. Consequently, we focus on RNAs with lengths between 16 and 256 nucleotides.

**Figure 5.**
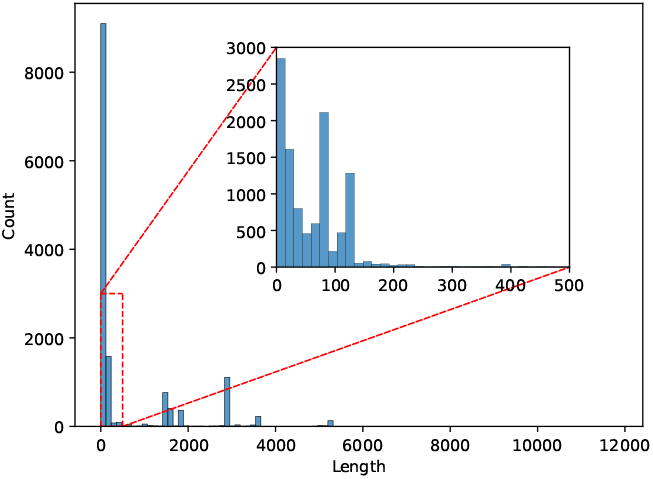
Sequence length distribution.

The distribution of MStA sizes for the filtered structures is depicted in Figure 6, with a concentration at both ends. Notably, 97% of the RNAs have a corresponding MStA (with MStA size *≥*1), and approximately half of these structures contain more than one thousand MStA elements. This results in a disproportionately large cluster, comprising 5,976 structures, formed during the clustering process. This cluster is included in the training set.

**Figure 6.**
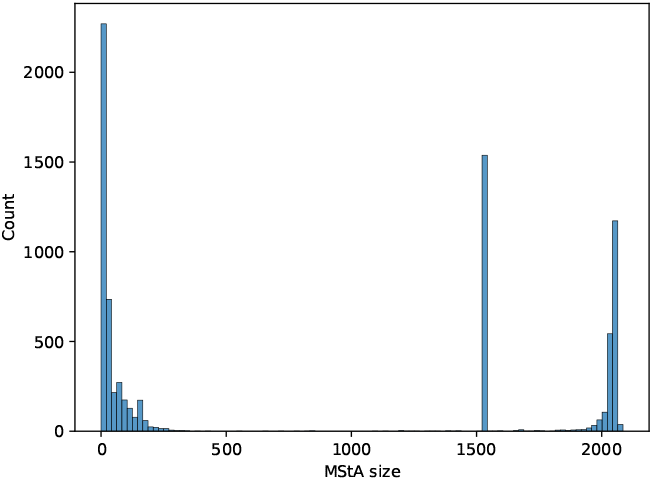
Size distribution of MStA in filtered structures.

After splitting, the sequence length distribution across the training, validation, and test sets is shown in Figure 7. All three sets encompass the full range of sequence lengths.

**Figure 7.**
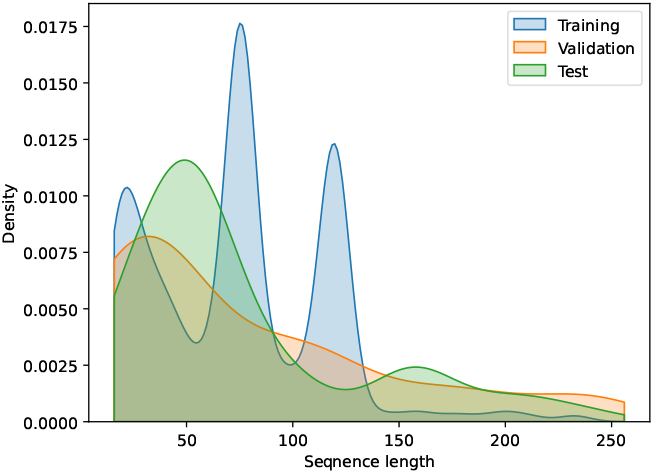
Sequence length distribution in training, validation, and test sets.

Finally, the joint distribution of sequence length and MStA size in the test set is presented in Figure 8. MStA structures are present for sequences of all lengths, and 10% of the test set contains no MStA, enabling a comprehensive evaluation of AlignIF under diverse scenarios.

**Figure 8.**
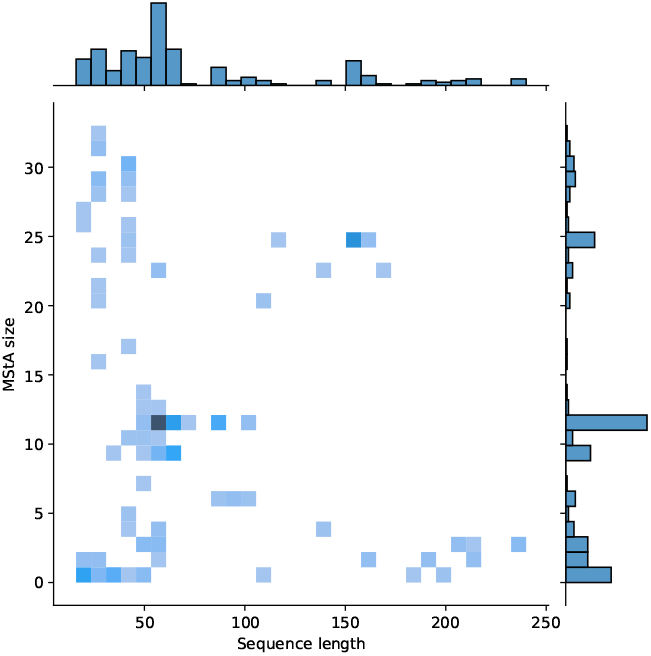
Bivariate distribution of sequence length versus MStA size in the test set.

### Local coordinate systems modified in our work

The local coordinate system traditionally developed to protein studies (Ingraham et al., 2019; Gao et al., 2022) and recently adapted for RNA (Tan et al., 2024).

In RDesign (Tan et al., 2024), the local coordinate system is defined using three consecutive phosphorus (P) atoms. Due to the flexibility of RNA chains, the local system in RDesign is highly dynamic, making it difficult to capture consistent features with rotation invariance. Furthermore, establishing the coordinate system at the beginning or end of the sequence is not feasible, as three successive residues are required for its definition.

In our work, we introduce a modified local coordinate system based on the position of atoms C2’, C3’, and C4’ within the solid sugar ring of each nucleotide residue. The system is described as follows:

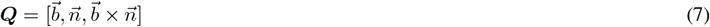

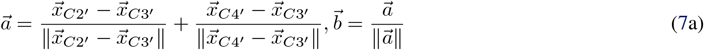

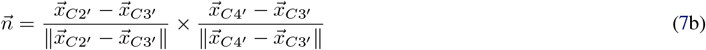

where 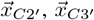, and 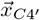 represent the coordinates of atoms C2’, C3’, C4’, respectively. 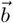 is the unit vector along the bisector of the angle formed by the rays of 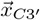 to 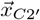 and 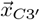, to (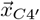 while 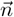 is the unit vector normal to the plane formed by these atoms. The modified system addresses the issues present in the previous system and achieves better performance.

